# *In Silico* Study of the Early Stages of Aggregation of *β*-Sheet Forming Antimicrobial Peptide GL13K

**DOI:** 10.1101/2024.01.25.577308

**Authors:** Mohammadreza Niknam Hamidabad, Natalya A. Watson, Lindsay N. Wright, R.A. Mansbach

## Abstract

Antimicrobial peptides (AMPs) are of growing interest as potential candidates for antibiotics to which antimicrobial resistance increases slowly. In this article, we perform the first *in silico* study of the synthetic *β* sheet-forming AMP GL13K. Through atomistic simulations of single and multipeptide systems under different charge conditions, we are able to shine a light on the short timescales of early aggregation. We find that isolated peptide conformations are primarily dictated by sequence rather than charge, whereas changing charge has a significant impact on the conformational free energy landscape of multi-peptide systems. We demonstrate that the lack of charge-charge repulsion is a sufficient minimal model for experimentally observed aggregation. Overall, our work explores the molecular biophysical underpinnings of the first stages of aggregation of a unique AMP, laying necessary groundwork for its further development as an antibiotic candidate.

## 1. Introduction

In the past decade, antimicrobial resistance (AMR), or a lack of response to traditional antibiotics, has been on the rise. In response, major health organizations such as the World Health Organization (WHO), have declared it one of the top ten threats to global health ^[1]^. At the same time, the drug delivery industry has been slow to deliver new antibiotics, especially those with new mechanisms of action, or those to which the build-up of AMR is expected to be comparatively slow. One reason for the difficulty in developing drugs with more complex modes of action is a lack of understanding of the molecular-level mechanisms contributing to various stages of activity in bacteria.

A good source of potential new antibiotics to which a comparatively slow build-up of resistance is observed are antimicrobial peptides (AMPs) ^[2]^. AMPs are short proteins that form an essential part of the nonspecific immune system in many organisms. Most are positively charged, which confers on them their ability to strongly interact with negatively-charged prokaryotic cell membranes in comparison with the neutral membranes of eukaryotic cells. AMPs are also typically amphipathic, which enhances their ability to interact as well with the strongly hydrophobic interior of the bacterial cell membrane bilayer ^[3]^. Many of them act against bacteria through disrupting the bacterial membranes, and many types of mechanisms have been proposed and reported in the literature ^[4,5]^. The most commonly studied AMPs are amphiphilic helix-formers that act via pore formation^[6–12]^.

A less well-studied group of AMPs is those that form *β* sheet-like structures at the bacterial membrane surface. The peptide GL13K (Fig. 1) is one of these and is believed to act through a carpet-like mechanism, involving micellization of the membrane and possibly the formation of transient pores ^[13]^. It has demonstrated promising applications to the development of antimicrobial surfaces ^[14–16]^, as well as the ability to bind the bacterial endotoxin LPS, in addition to its membrane-disruptive ability. GL13K was derived from the Human salivary protein (BPIFA2), a lipopolysaccharide-binding protein that is considered to be an oral host-defense protein. It was created by isolating residues 141 to 153 in this peptide (named GL13NH2) and replacing the glutamine, asparagine, and aspartic acid residues in positions 2, 5, and 11 with lysines ^[17]^.

**Figure 1.**
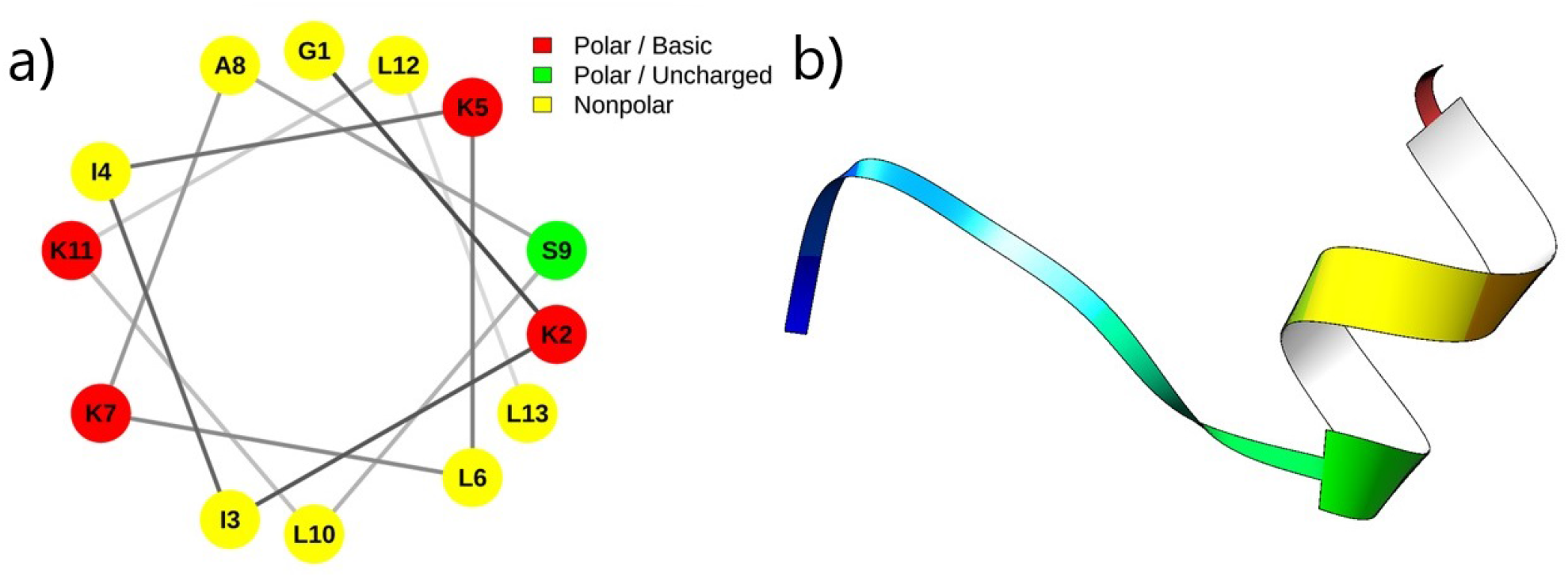
GL13K sequence and initial modeling structure. (a) A helical wheel representation of GL13K sequence is depicted, created using the NetWheels interface ^[18]^. The sequence contains four charged (basic) residues, eight nonpolar residues, and one polar residue (SER9). (b) Initial secondary structure of GL13K predicted by PEP-FOLD 3 web-server ^[19]^ is visualized by Chimera package^[20]^.

GL13K has been well-characterized experimentally on long time-scales. It is known to interact with both bacterial membranes in general and the bacterial endotoxin LPS in specific. Near bacterial-like membranes and in solution at high pH, it undergoes aggregation into large non-amyloidogenic aggregates with high *β* sheet content on time-scales of days, but near eukaryotic membranes and in solution at closer to neutral pH it is primarily unstructured in solution. Modification of the sequence, particularly replacement of any of the positively-charged lysine residues, has a significant impact on the rate of aggregation and the observed morphology of the resulting aggregates and may even prevent self-assembly entirely ^[13,17,21–24]^.

Despite the comparatively numerous experimental studies that have been performed on GL13K, there have been no computational studies, and understanding of the molecular-level drivers of single-peptide conformations and the early stages of aggregation remains limited. In this work, we employ atomistic molecular dynamics to characterize single-peptide conformations and early-stage aggregation of GL13K. We investigate the role of charge, sequence, and multi-body effects and demonstrate that the mechanism of aggregation under high pH conditions can be explained primarily through loss of charge-charge repulsion between peptide monomers. In the first part of the results and discussion section (Sec. 2.1), we discuss simulations of isolated peptides in water and probe the effects of charge and sequence via assessment of the secondary structure, radius of gyration, and contact maps. Next, we simulate charged and uncharged systems of eight GL13K peptides and assess the properties of early-stage aggregation through comparison of cluster size and *β* sheet content over time (Sec. 2.2). Finally, we compare the conformational landscapes of single and multi-peptide GL13K via hydrogen-bonding and free energy surface analysis (Sec. 2.3). Overall our work provides an initial microscopic analysis of the detailed molecular-level contributions to early-stage assembly of GL13K.

## 2. Results and Discussion

We studied seven different single-peptide systems and two different multi-peptide systems to understand the effects of sequence and charge on peptide conformation and early-stage aggregation. The sequences of the single-peptide systems studied are shown in Table 1, with changes from the original GL13K system highlighted in red: they include three alanine substitution variants previously studied experimentally ^[25]^; GL13K with its lysines charged, to model its behavior in a neutral pH; GL13K-neutral with its lysines deprotonated (uncharged), to model its behavior in a high pH environment; GL13K-R1, a peptide with the same amino acid composition as GL13K but a different sequence ^[21]^; GL13K-R1-neutral, employing the R1 sequence with its lysines deprotonated. GL13K-R1 functions as a demonstration of the importance of specific sequence, rather than solely composition as well as a validation of the computational model. We also use it to assess the validity of the computational model.

**Table 1.**
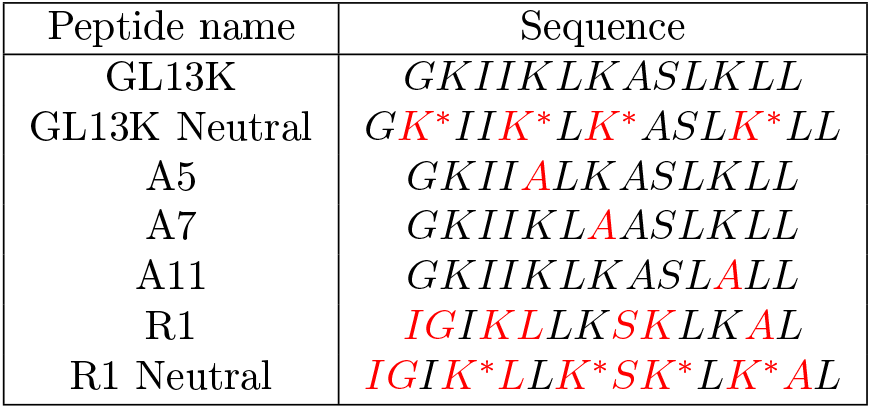
Systems studied: list of amino acid sequences for all different systems studied in this article, with naming conventions indicated on the left, sequence differences from the original GL13K highlighted in red, and asterisk indicating deprotonation of an amino acid.

Although lysine is deprotonated at a sufficiently high pH, to more accurately assess the direct effect of changing pH on a molecular system, it is also possible to employ techniques such as constant-pH molecular dynamics ^[26]^. We do not do so for several reasons. Firstly, this allows us to directly test a simpler hypothesis: that observed aggregation and secondary structure change can result primarily from the amount of average charge on the peptide. Second, for the more computationally intensive simulations of aggregation, it allows us to ameliorate the requisite computational resources, and we have previously demonstrated that although it is a simple approximation, it is sufficient to model the effects of pH on aggregation in other systems of aggregating short peptides ^[27]^.

### 2.1 Charge and sequence effects in single-peptide systems are responsible for alpha helices but not beta sheets

To understand the conformation and the shape of a single peptide in solution, we calculated shape parameters and contact maps. Using the MDTraj scientific package. ^[28]^, we computed the radius of gyration (*R*_*g*_), acylindricity (*a*_*cyl*_), and asphericity(*a*_*sph*_), which are all quantities derived from the principal moments 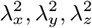of the gyration tensor of the peptide ^[29]^.

Radius of gyration is defined as,

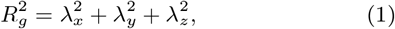

and it is a measure of how extended or compact the peptide is.

Asphericity is calculated as,

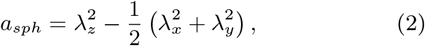

and it is a measure of how symmetrically the peptides’ residues are positioned about all three axes. A larger value of asphericity corresponds to a less spherical or less symmetric arrangement, while a value of zero corresponds to spherical symmetry or similar.

Acylindricity is calculated as,

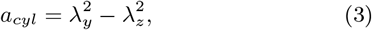

and it is similarly a measure of the deviation from cylindrical symmetry, with a zero value corresponding to equal values of the two shorter coordinate axes.

In Fig. 2, we illustrate the (a) the per-system average radius of gyration *⟨Rg⟩*, (b) per-system average asphericity *⟨a*_sph_ *⟩*, and (c) per-system average acylindricity *⟨a*_cyl_*⟩* . We observe that the average radius of gyration (a) ranges from 1.02 *±* 0.02 nm for GL13K to 0.74 *±* 0.01 nm for R1neutral and we note significant differences between the *R*_*g*_ of different systems (Anova test, significance level=0.05 and p-value=1.37E-08). In addition, the two neutral systems demonstrate overall lower gyration radii than their charged counterparts visually, though only the difference between GL13K and GL13K-neutral is significant (*t*-test, significance level=0.05 and *p*-value=1.12E-02). As might be expected, greater charge-charge repulsion leads to more extended conformations. Finally, we note that the A7 (0.98 *±* 0.03 nm) and A11 (0.98 *±* 0.01 nm) variants have the most similar gyration radii to the GL13K system, whereas the A5 variant (0.90 *±* 0.03 nm) tends to be somewhat more collapsed. We note similar trends in the asphericity of the system, where systems with higher radii of gyration tend also to be more aspherical, as their conformations tend to be more extended and thus one moment of the gyration tensor is longer than the other two. Asphericity values range from 0.30 *±* 0.05 nm, for the most spherical R1 and 0.24 *±* 0.01 nm for R1-neutral systems, to 0.58 *±* 0.02 nm for the GL13Kneutral and 0.51 *±* 0.05 nm for A5 systems, to nearly 0.75 nm (0.74 *±* 0.04 nm for GL13K, 0.70 *±* 0.04 nm for A7, and 0.66 *±* 0.01 nm for A11). Finally, acylindricities tend to be rather low across the board, suggesting all peptides take on comparatively cylindrically symmetric shapes.

**Figure 2.**
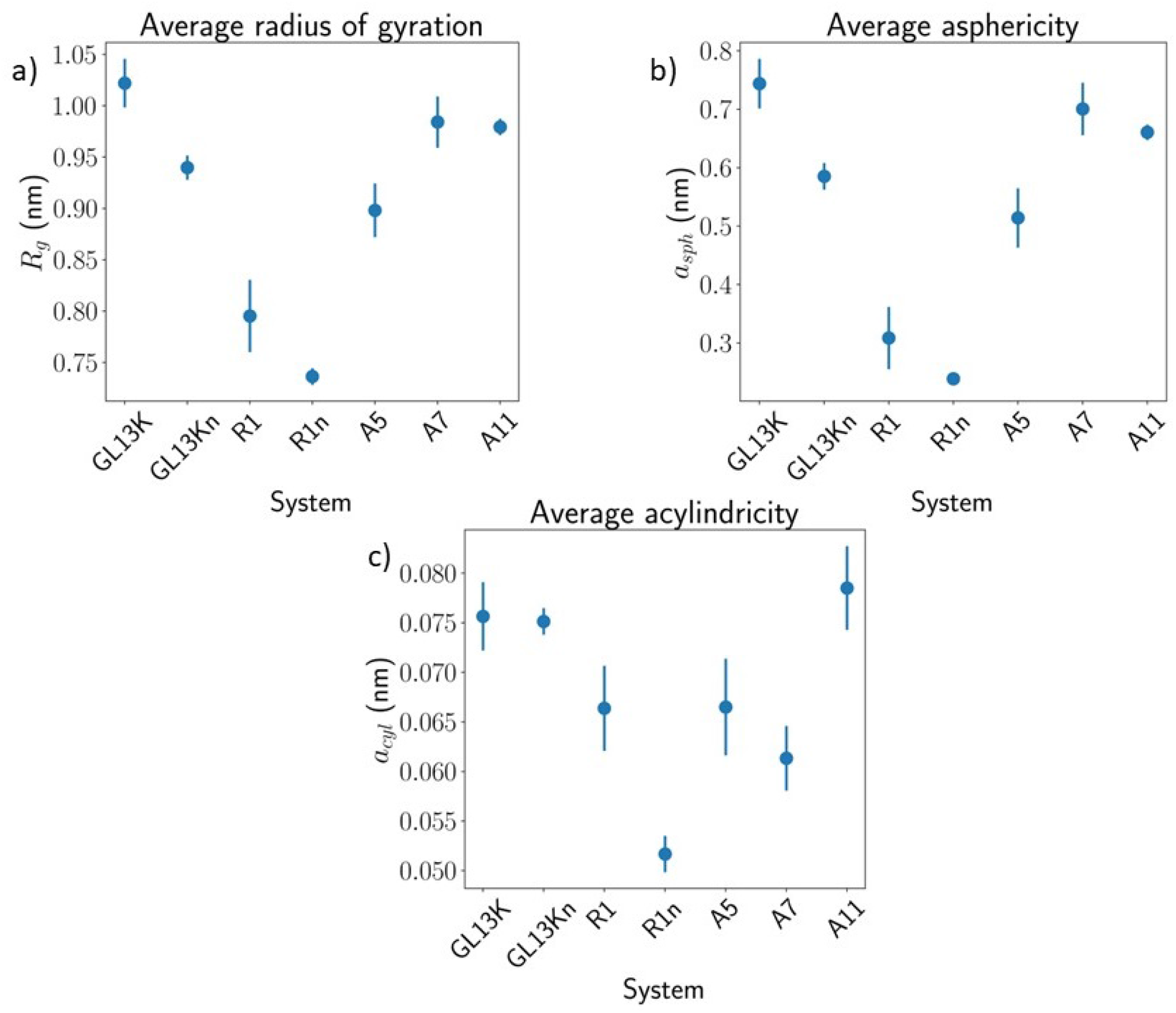
Average shape parameters of different single-peptide systems. (a) Radius of gyration *R*_*g*_, (b) asphericity *a*_sph_, (c) acylindricity *a*_cyl_. Averages are taken across the final 400 ns of five different replicas. Standard error indicates the standard error in the means of the five different replicas.

To gain a greater understanding of the individual residue contributions to the changing shapes of the system, we also investigated the average per-residue contact maps (Fig. 3) demonstrating the intramolecular residue-residue distances. GL13K and its A7 and A11 variant are the most extended, as also measured by the radius of gyration. On average, they also demonstrate a linear increase in the physical distances of residues that are further apart in the sequence, up to a maximum end-to-end distance of 2.5 *±* 0.1 nm. GL13Kneutral demonstrates a very similar contact map, with the same linear increase, but up to a smaller maximum end-to-end distance of 1.6 *±* 0.1 nm. R1 and R1-neutral are far more compact and demonstrate equidistant blocks (which in the next paragraph we associate with residues equidistant due to being on the same helical twist); R1-neutral has a longer such block. Finally, A5 is the most collapsed, with most of its C-terminus being roughly the same distance from the Nterminus, concomitant with its lower *R*_*g*_ and corresponding to the repeated sampling of a folded-over *β*-hairpin-like configuration. Evidently, the mutation of LYS5 in ALA results in collapse of the N-terminal tail over the peptide, whereas this does not occur when all lysines are neutralized, suggesting that either ALA may play a role in stabilizing this conformation or that charge asymmetry does.

**Figure 3.**
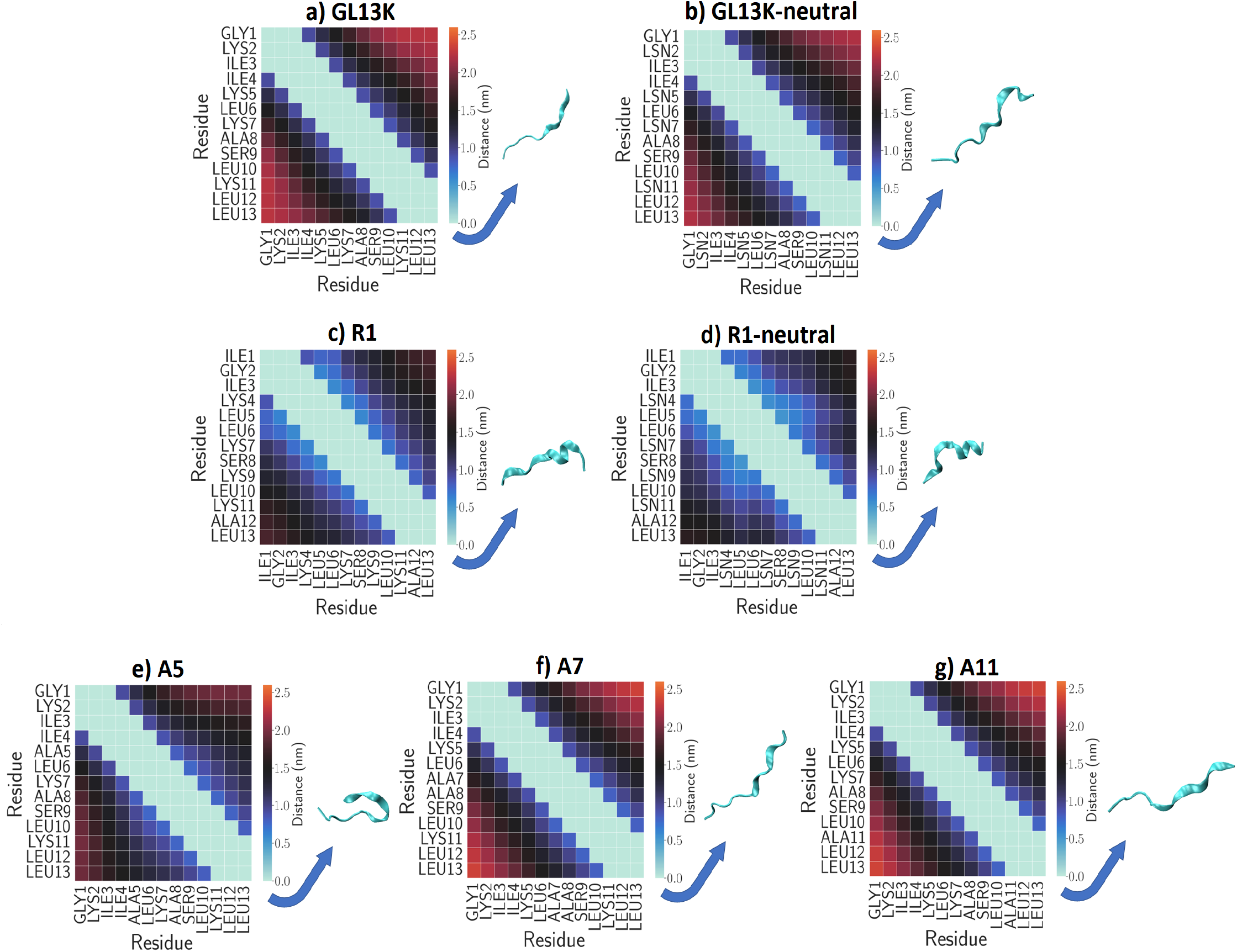
Average contact map of single-peptide systems in solution for (a) GL13K, (b) GL13K-neutral, (c) R1, (d) R1-neutral (e) A5, (f) A7, (g) A11. Averages are taken across five independent replicas, and the standard error is shown in Fig. S3 . Representative conformations of each peptide were rendered in VMD^[30]^ and displayed to the right hand side of their respective variant’s contact map.

To investigate the effect of charge and peptide sequence on the secondary structure of the peptide, we calculated the per-residue average secondary structure over all time, using compute dssp tool of MDTraj package ^[28]^. We plot average secondary structure and its error in Fig. 4 and Fig. S1 respectively. (In Fig. S2 in the Supplementary Information, we also show the average secondary structure of each peptide over time, averaged across all residues.) We observe in GL13K (Fig.4a) primarily extended-loop conformations of all residues (92.92 *±* 4.05%), with some slight (0.14 *±* 0.03 of the time) bend and turn primarily localized around LYS7, ALA8, and SER9. ILE4, LYS5, and LEU6 and LEU10, LYS11, LEU12 all demonstrate a small amount *β*-ladder content, probably corresponding to a rare folded-over conformation with short intra-molecular *β* sheets (cf. conformation illustrated in Fig. 9a, bottom left). Overall, however, GL13K primarily retains an extended structure with all residues in random coil conformations. GL13K-neutral (Fig. 4b) likewise demonstrates primarily extended random-coil structure for most of the residues. There is a slightly lower preponderance towards random coil for ALA8, SER9, and LYS11 ( 46.2 *±* 4.3%, 51.7 *±* 3.9%, and 47.7 *±* 2.7%, respectively), with a concomitant marginal increase in *α*helical content centered on these three residues, but overall both GL13K and GL13K-neutral sample random-coil conformations the bulk of the time, demonstrating little change in their secondary structure with charge neutralization. The A5, A7, and A11 variants (Fig. 4 e-g, all charged), sample similar configurational space. A5 has a slight enhancement in helix and turn content, centered around ALA8, presumably due to its preference for the *β*-hairpin-like configuration. In comparison, R1 and R1-neutral (Fig. 4c-d) both demonstrate comparatively extended helical structures from residues *∼*3 *−*10 (R1) and residues *∼*2 *−*10 (R1-neutral), with R1-neutral displaying heightened helical content in the N-terminal half and slightly heightened turn content near SER8.

**Figure 4.**
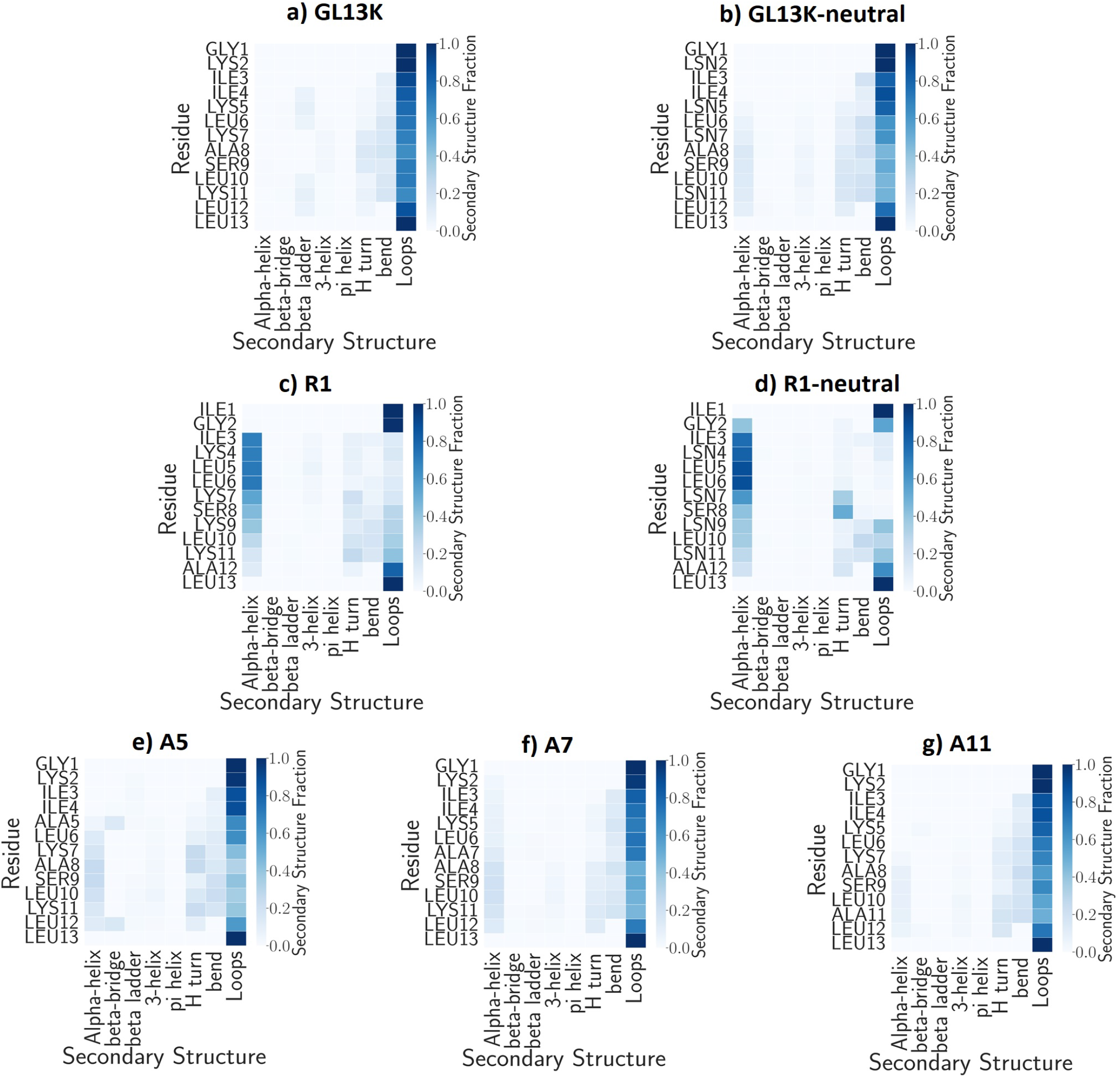
Average secondary structure of single-peptide systems in solution

Our simulation results are generally in accord with experimental observations, providing *a posteriori* evidence for our modelling procedure. In addition, there are some points of interest that we can use to understand molecular-level origins of *β*-sheet aggregation. Previous circular dichroism experiments ^[21]^ under changing pH conditions reported some *α*-helical content in GL13K, which we do not observe at our proxy neutral pH; however, their experiments start at pH 8.5. Additionally, they observed *∼* 25% *β*-sheet content at pH 8.5, increasing to *∼* 45% at pH 10.6, whereas we observe *<* 10% *β* bridge content in GL13K and none in GL13K-neutral. They did, however, observe *α*-helical content in R1, increasing with increasing pH (lowered charge), as do we. We also observe that GL13K at pH 7 takes on primarily random-coil structure as expected^[21,31]^. Apart from some slight collapse, likely due to the loss of charge repulsion in the system, we do not observe significant conformational change when the lysines are neutralized, unlike what is observed experimentally. Thus, we demonstrate that the preponderance of observed *β* sheet content arises from inter-molecular peptide interactions, with potentially some slight contributions from intra-molecular interactions between residues 4-6 and residues 10-12 at pH levels close to neutral.

Finally, it was observed that beginning at pH 9.5, where we expect fewer charges on the lysines, aggregation took place into twisted *β*-sheets in the GL13K and A7 systems, with slower formation of similar structures in A5 and more helical content in aggregates in A11 ^[25]^. Our results agree in the sense that A7 behaves similarly to GL13K at pH 7, suggesting that the lysine at position 7 does not strongly impact its conformation one way or the other. Presumably, the removal of the charge at A5 is responsible for its greater collapse, since residues 5 and 11 will no longer repel one another. Apparently, the repulsion between LYS7 and LYS11 is sufficient to keep A11 in a somewhat more extended conformation.

### 2.2 Peptide aggregation is seeded by multimeric dynamic equilibrium in the charged state

In this section, we investigate the aggregation behavior of GL13K by studying eight peptides in the charged (protonated, proxy for neutral pH) and uncharged (deprotonated, proxy for high pH) states. To understand the general behavior of multi-peptide systems, we have plotted the average system cluster size *⟨ N*_ave_*⟩* (average taken across all clusters across all replicas) and average maximum cluster size *⟨ N*_max_ *⟩*(average of maximum cluster size in each replica) over time for both uncharged and charged peptides (Fig. 5). Two peptides are defined to be in the same cluster if the distance between two non-hydrogen atoms from different peptides is less than or equal to a cut-off distance. We define our cutoff distance to be 5Å, as has been used previously, because this value is close to the minimum of the Lennard-Jones potential ^[32–35]^. We observe aggregate formation in both charged and uncharged systems, but aggregation proceeds to a greater extent in the uncharged system, presumably due to repulsion of the lysine residues in the charged system. In the charged system, the average cluster size remains close to monomeric (1.5 *±* 0.2), but semi-persistent dimers and transient clusters up to size six are observed (cf. 5a,c). The overall system appears to rapidly attain a dynamic equilibrium. The uncharged system undergoes non-equilibrium irreversible aggregation to the maximum possible size of eight peptides (cf. 5a,b,d) in a cluster on a timescale of 431 *±* 28 nanoseconds, with error estimated from five independent replicas.

**Figure 5.**
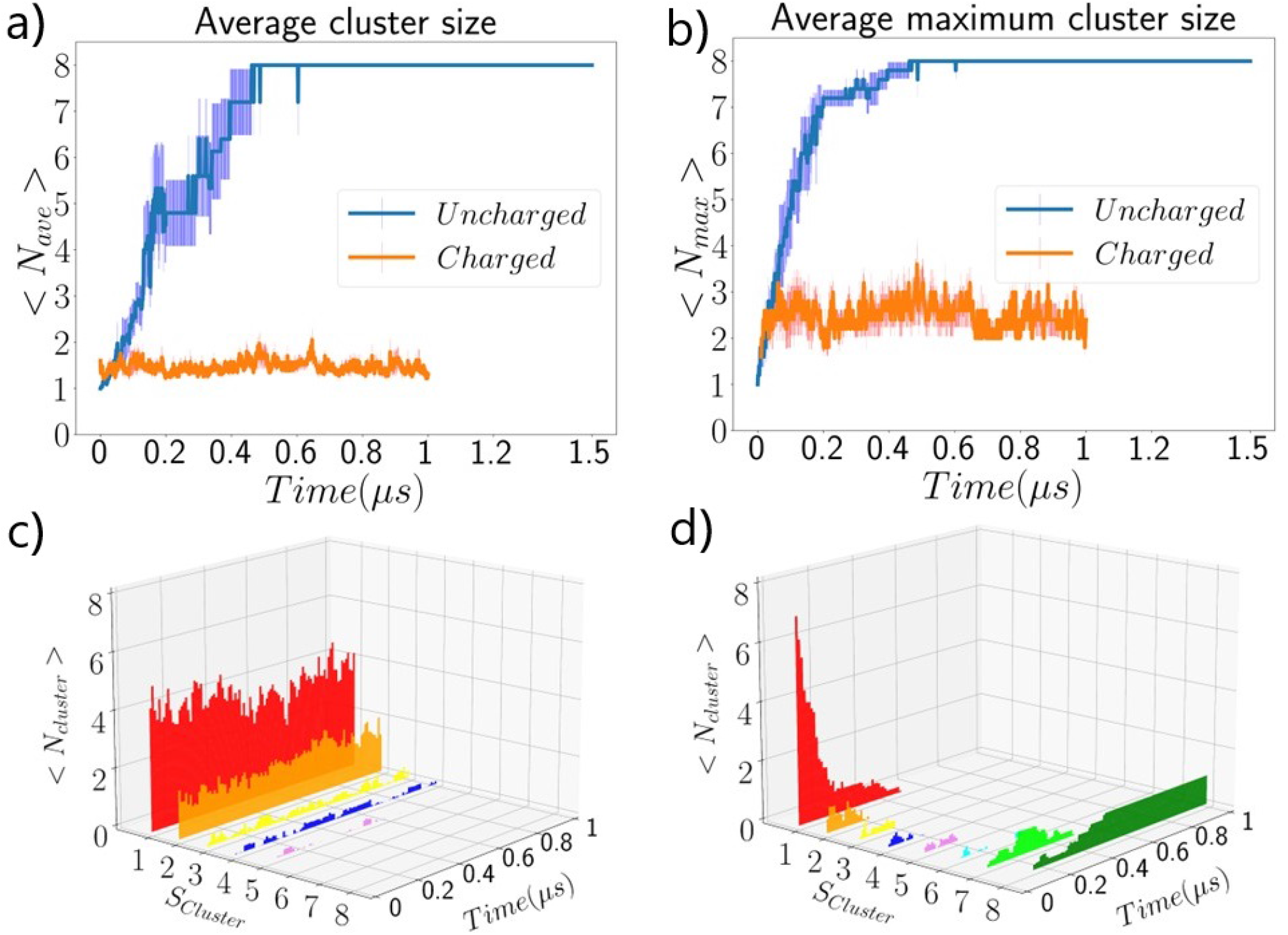
Growth of clusters in multi-peptide systems. (a) Average cluster size *N*_ave_ versus time. (b) Maximum cluster size *N*_max_ versus time. For (a)-(b) we display the average and standard error across five independent replicas for both charged and uncharged peptides. (c)-(d) Sampling of clusters of individual sizes *S*_*Cluster*_ =1–8 with time for (c) charged and (d) uncharged systems.

In Fig. 6, we present the average *β* sheet content of the multi-peptide systems with respect to time and cluster size. *β* sheet content was calculated by using the DSSP package from the mdtraj Python package ^[28]^ to identify the secondary structure of each residue at each timestep according to backbone angles and sidechain positions. Due to the evidence of non-equilibrium afforded by a changing radius of gyration, the first 100 ns simulation time of both charged and uncharged systems was not considered for this calculation. At each timestep, the fraction of *β* sheet content *β*_frac_ was defined as the number of residues out of the 8 *×* 13 = 104 total in which the ‘E’ identifier appeared. We observe an increase over time of *β* sheet content for both charged and uncharged systems, and a consistently higher amount of *β* sheet content in the uncharged system. The uncharged system appears to reach an equilibrium value of 0.35 *±* 0.05 after 1100 ns of growth, while the charged system appears to reach an equilibrium value of 0.12 *±* 0.04 after 425 ns. Based on the relationship between cluster size and *β* sheet content, we note heightened *β* sheet content in the uncharged system for aggregates as small as size 3, but we also note an increase in the percentage of *β* sheet content with increasing aggregate size, up to about size 5, after which it levels off. In addition, only for clusters of size greater than 3 in the uncharged system is there significant *β*-sheet content in excess of 10%. Since, in the uncharged system, the maximum cluster size is attained on timescales of 500-600 ns, concomitant with an initial steep increase in *β*_frac_, followed by a slower continued increase in *β*_frac_, this suggests that aggregates form rapidly with a bulk of the *β* sheets formed on initial aggregation and rearrange on longer time-scales in ways that moderately increase *β* sheet content. This is also supported by the comparatively broad distributions of *β*_frac_ with respect to cluster size and by the initial increase in *β* sheet content in the uncharged system despite its rapidly attaining a dynamic equilibrium in terms of cluster formation.

**Figure 6.**
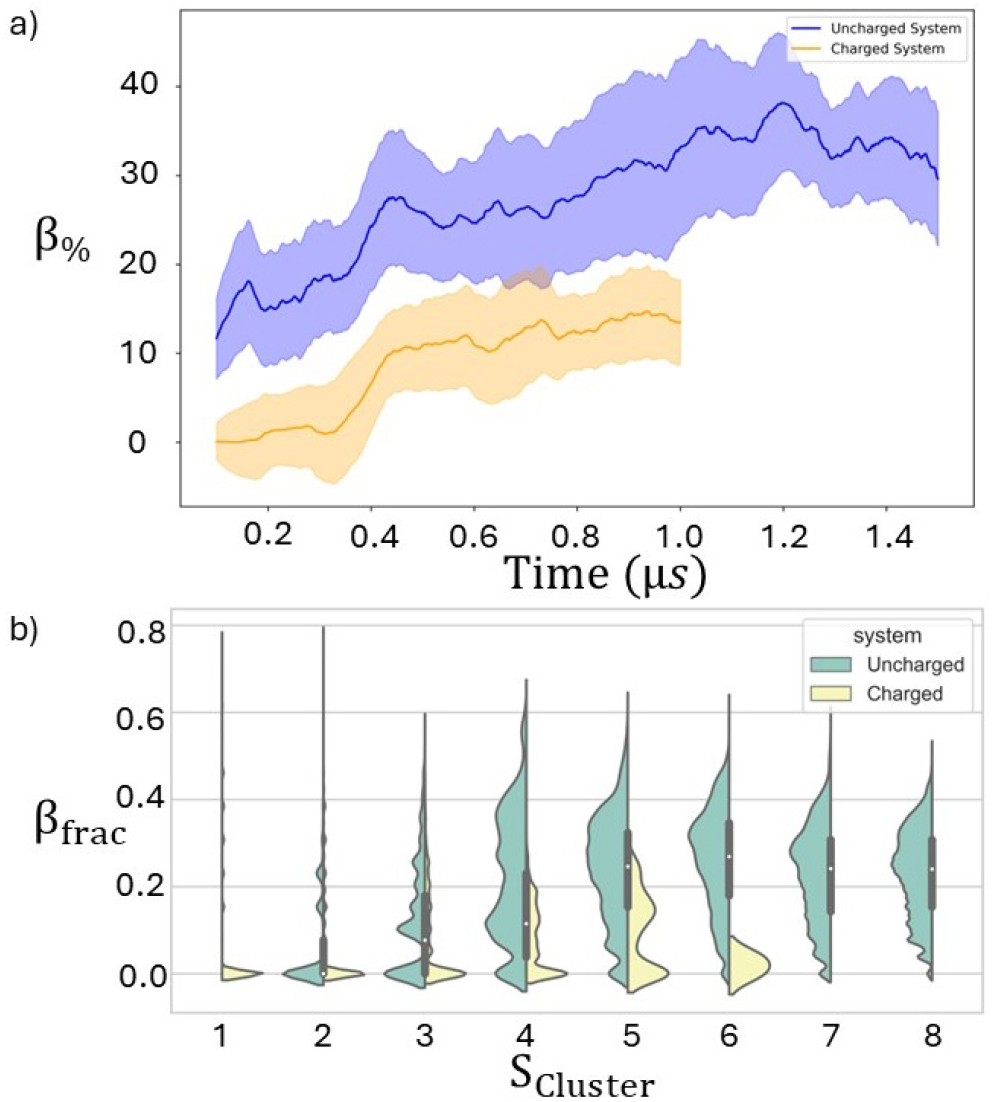
Average *β*-sheet content versus time of multi-peptide charged (in red) and uncharged (in blue) systems. We plot the fraction of residues in the system that demonstrate *β*-sheet content *β*_frac_ (a) versus time and (b) versus cluster size *S*_Cluster_. Distributions in (b) are normalized to the same maximum value. Averages and standard error are taken across five independent replicas. The uncharged system was run for 500 ns longer than the charged one to ensure it was equilibrated.

Qualitatively, our observations are consistent with experimental ones, which saw an increase from about 25% *β* sheet content at pH 8.5 to about 45% *β* sheet content at pH 10.6 ^[21]^. Quantitatively it is 5-10% lower at neutral pH and 5-10% lower at pH > 10.6. This, particularly the latter, suggests modest long-timescale rearrangements of largescale aggregates towards increasing *β* sheet content. Since our results already suggest longer time-scale aggregate relaxation and experimentally observed large-scale aggregate growth is quite slow (on the order of days), this hypothesis seems well-supported.

In Fig. 7, we illustrate the primary locations of *β* sheet formation via a per-residue secondary structure map, where the percentage of secondary structure is measured over the final 500 ns of all simulations under consideration, with standard error as computed across five replicates demonstrated in Fig S4. We observe primary formation of the *β* sheets in the aggregating, uncharged system near the Nterminus, particularly in residues 4-6, all of which have *∼* 40% *β* sheet content. Appreciable *β* sheet content is also found in residues 7-12, which contain 20*−* 30% *β* sheet content at equilibrium. The terminal residues largely do not participate in *β* sheet formation on the timescale of our simulations. The residues that demonstrate lower *β* sheet content (7-12) in the uncharged system are those demonstrating appreciable (roughly similar content) in the charged system, suggesting transient aggregates are stabilized by hydrogen-bonding near the C-terminal end, likely because of the lower density of charged residues. Thus the increase in *β* sheet content with increasing pH (decreasing charge) largely comes from enhanced interaction of the residues on the N-terminal side of the peptide.

**Figure 7.**
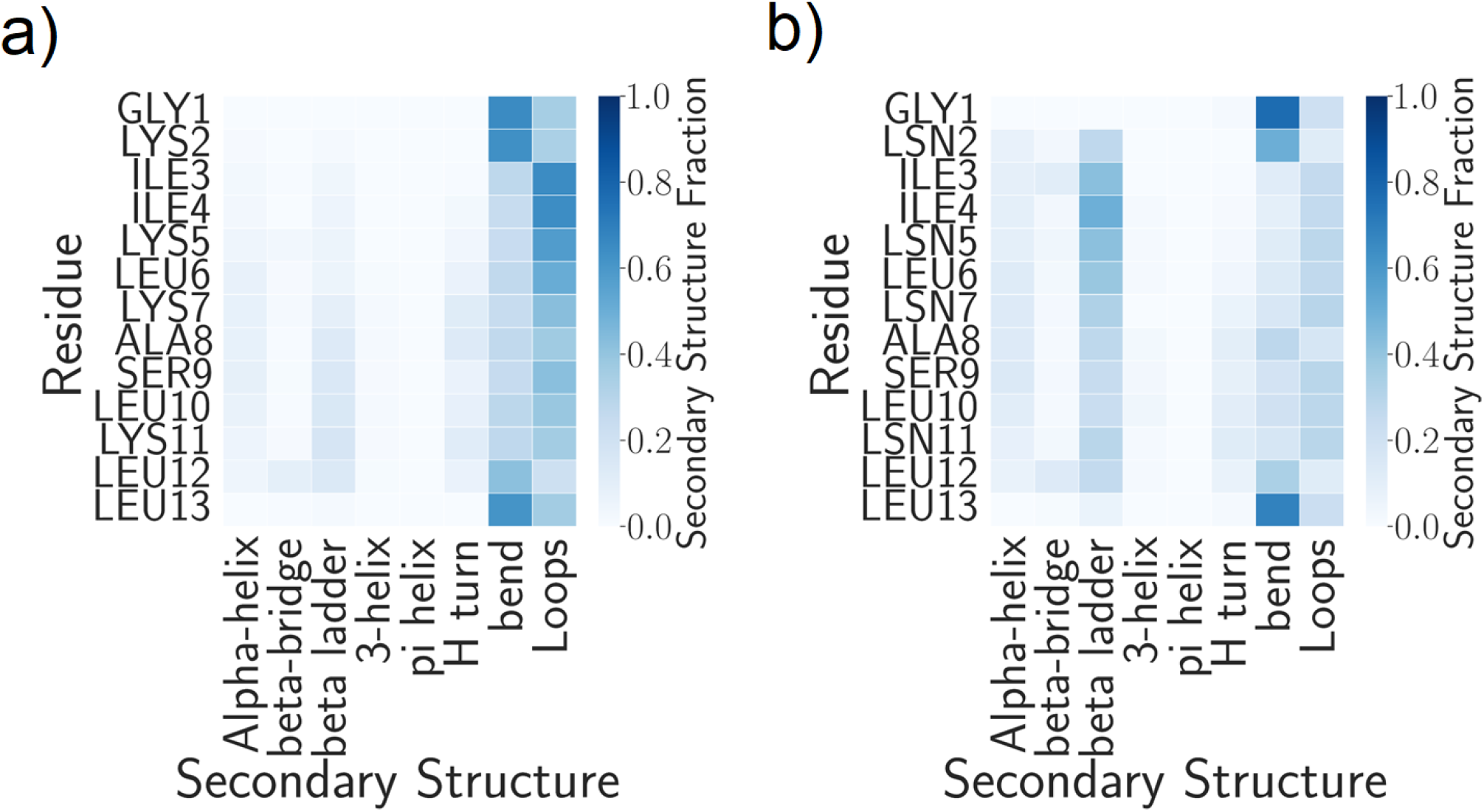
Average secondary structure per residue of multi-peptide systems: (a) charged system; (b) uncharged system. Averages are taken over the last 500 ns of all five simulations [after the *β*-sheet percentage has reached an equilibrium value (approximately 550 ns for charged, 1100 ns for uncharged systems), see Fig. 6]. Percent error is shown in Supplementary Figure S4.

### 2.3 Changes to conformational ensemble in single and multi-peptide systems

In this section, we focus on the different conformations taken on by single and multi-peptide GL13K in charged (proxy for neutral pH) and uncharged (proxy for high pH) conditions. We compare the systems that do and do not involve multibody interactions through investigating the locations and extent of hydrogen bonding and through visualization of the overall free energy landscapes.

In Fig. 8, we present an analysis of the hydrogen bonding networks of the charged and uncharged single and multipeptide systems. We compute h-bonds using the gmx hbond command of the Gromacs package with the following criteria defining an h-bond: Cutoff radius and Cutoff angle 0.35 nm and 30 degrees between Hydrogen-Donor-Acceptor, respectively. For single peptide systems, we show the average number of intra-molecular hydrogen bonds (h-bonds) that are present in at least 10% of the post-equilibration simulation time (Fig. 8a,d). For multi-peptide systems, we show the average number of intra-molecular and inter-molecular h-bonds that are present in at least 10% of the final 500 ns of simulation (Fig. 8b,e) and those that are present in at least 40% of the final 500 ns of simulation (Fig. 8c,f). We note that intra-molecular hydrogen bonds in the single peptide systems are quite transient and none of them are present in at least 40% of the simulation time. Standard errors are computed from the standard error of the means of five independent simulations.

**Figure 8.**
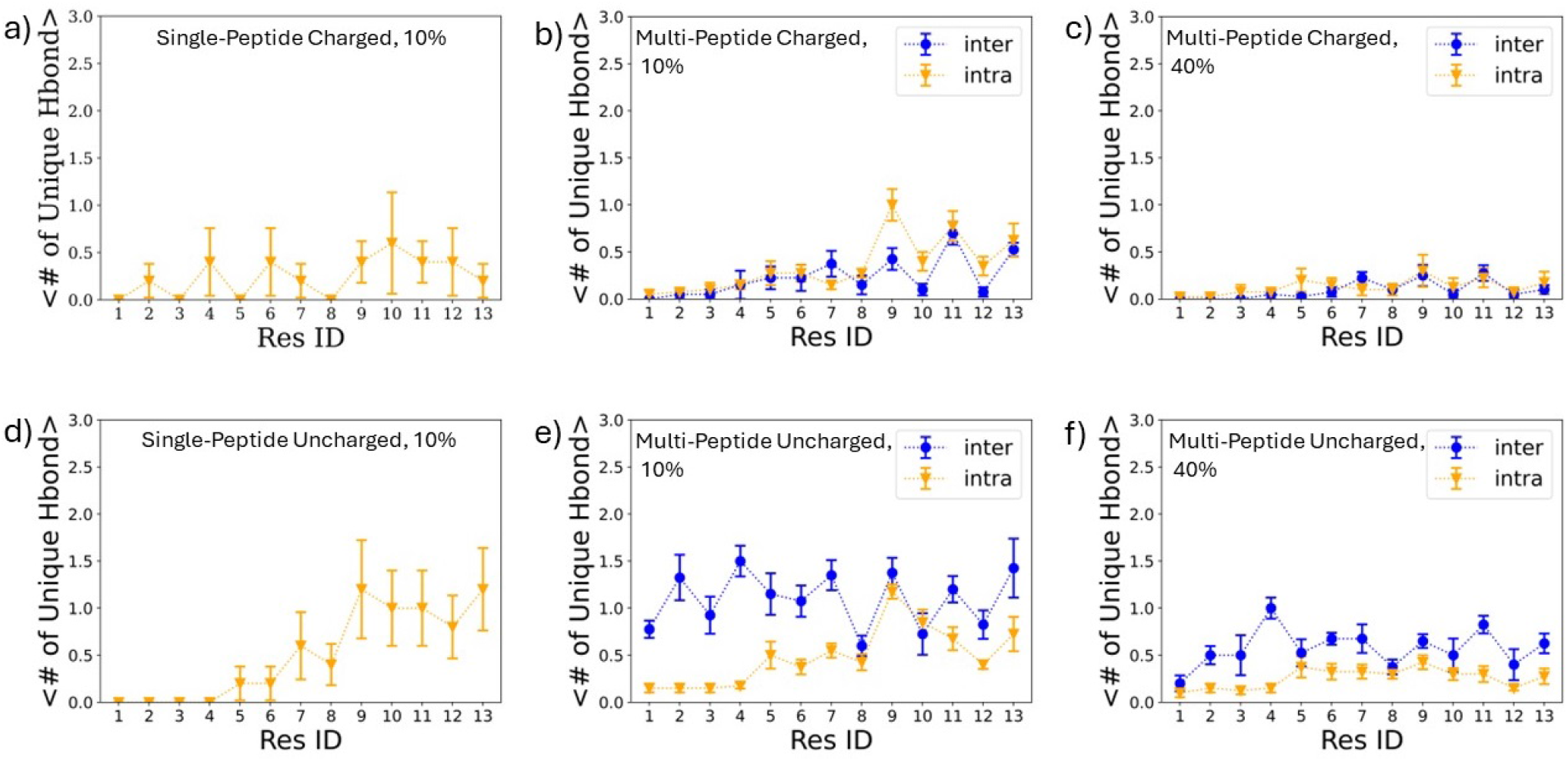
Hydrogen bonding patterns for single and multi-peptide systems. a),d) Single peptide systems, a) charged, d) uncharged. Average number of h-bonds per residue found in at least 10% of the final 400 ns of simulation time. b-c),e-f) Multi-peptide systems, b-c) charged, e-f) uncharged. b),e) depict the average number of h-bonds per residue found in at least 10% of the final 500 ns of simulation time. c),f) depict the average number of h-bonds per residue found in at least 40% of the final 500 ns of simulation time. Averages and standard error are taken over five independent replicas.

We observe in single-peptide systems an elevation in transient h-bonds, particularly in residues 7-13 on the C-terminal side of the peptide for the uncharged versus the charged system. There are few even transient h-bonds in the charged system, presumably due to electrostatic repulsion, but there is some small h-bonding present on the N-terminal end, for residues LYS2, and ILE4. Particularly notable increases (from *<* 1 in the charged system to almost 2 h-bonds that form transiently in the uncharged system) occur for SER9 and LEU13, the C-terminus. Of note, it is residues 4-6 and 10-12 that participate in the *β* ladder occasionally observed in the charged system (cf. Fig. 4), and ILE4 is one of the only residues to show higher h-bonding in the charged system over the uncharged one.

In the multi-peptide simulations, transient (*>* 10% formation) intra-molecular h-bonds are still observed for SER9 and LEU13, but there is no significant change between the charged and uncharged systems in this case. ILE4 participates in a very small amount of transient h-bonding in the charged system, but the primary intramolecular h-bonding near residues 4-6 occurs for 5-6 in the uncharged system. Overall, the largest pattern in intra-molecular h-bonding is a small increase of semi-persistent (*>* 40% formation) for residues 5-13. We note also, as expected, an increase in both transient (*>* 10% lifetime) and semi-persistent (*>* 40% lifetime) inter-molecular h-bond interactions in the uncharged system compared to the charged one. A significant increase in inter-molecular h-bonding near the N-terminal end for the uncharged system is consistent with the formation of *β* sheet aggregates stabilized on that side. The most persistent h-bond is formed for ILE4. Residues 9 and 11 demonstrate a larger increase in semi-persistent than in transient h-bonding patterns, corresponding to the observed formation of transient aggregates with minor *β*-sheet content near the C-terminus in the charged system. SER9, in specific, is probably the residue that most consistently, regardless of charge, participates in both intra-molecular and intermolecular hydrogen bonding, though the inter-molecular hydrogen bonding in which it participates is more persistent.

As a final step to investigate the overall impact of aggregation on the system, in Fig. 9, we compute free energy changes of single peptide and multi-peptide systems of charged and uncharged GL13K as a function of the first and second moments of the gyration tensor *G*_1_, *G*_2_ and the endto-end distance *R*_*e*2*e*_. We represent these computations as two-dimensional free energy surfaces (FESes). Although it is possible to perform more sophisticated analyses to identify relevant (typically slowest-evolving) shared collective variables (CVs) in which to construct FESes, we do not do so here. It has been previously demonstrated that for short, partially-disordered peptides, the three variables we have chosen are commonly related to the slower CVs of the system. In addition, from the previous analysis in this article, we believe that the ease of interpretation, in this case, outweighs the need to identify the single slowest or most representative variables. Finally, the identification of unified multi-system CVs–in other words, the CVs most broadly relevant to multiple systems under different conditions such as these–is still an area of active research ^[36–40]^.

**Figure 9.**
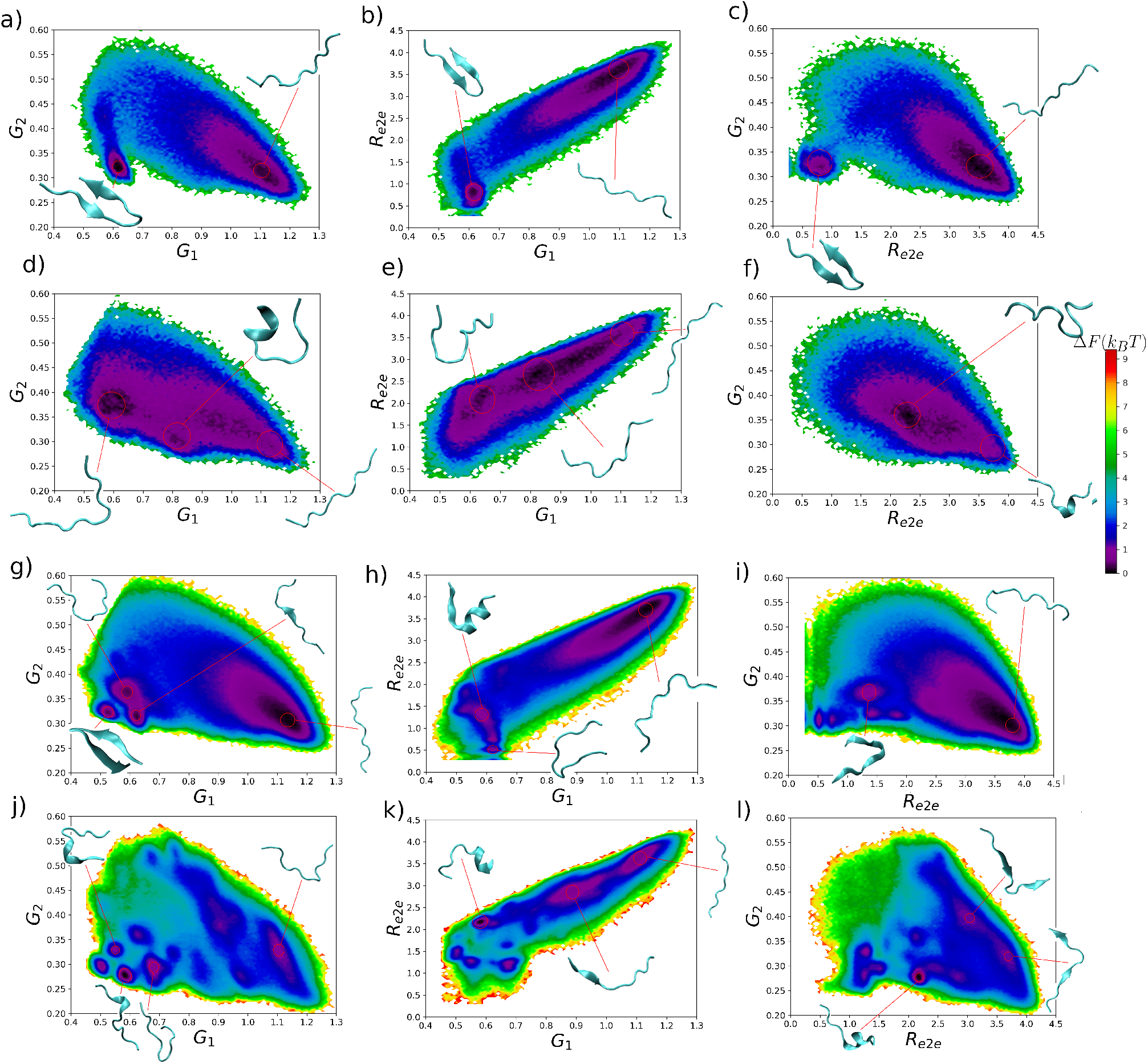
Comparison of two-dimensional free energy surfaces of (a-c) single peptide charged systems, (d-f) single peptide uncharged systems, (g-i) multi-peptide charged systems, and (j-l) multi-peptide uncharged systems. We choose to project into three collective variables known to be relevant to the systems of short, partially disordered peptides: the end-to-end distance in nanometers (*R*_e2e_), the 1st, and 2nd moment of the gyration tensor *G*_1_*andG*_2_. Free energy surfaces are computed over the final 400 ns of simulation for the single-peptide systems and for the final 500 ns of simulation for the multi-peptide systems. Surfaces constructed using the PyEmma package’s plot_free_energy function ^[41]^.

Free energy surfaces are computed and visualized using the `plot_free_energy` function of the PyEmma package ^[41]^ with the lowest and highest free energy values of 0.0 and 10.0 *k*_*b*_*T* respectively, meaning that the minimum is set arbitrarily to zero for each separate computation. Free energy is computed as Δ*F* = *k*_*b*_*T* ln *P*, where *k*_*b*_ is Boltzmann’s constant, *T* is the temperature of the simulation, and *P* is the probability of the relevant macrostate typically approximated as the number of times a certain set of values are visited within a simulation divided by the total number of timesteps. We also indicate on the free energy plots representative conformations from visually-identified free energy basins. For the single-peptide systems, FESes are computed from the single-peptide conformations of the last 400 ns. For the multi-peptide systems, FESes are computed from all single-peptide conformations from the last 500 ns. This results in slightly over eight times as much data for the multi-peptide systems, but the multi-peptide ones are still assessed in terms of single-peptide conformational ensembles.

For the single-peptide system, we observe that the charged system has two distinct free energy basins separated by *∼* 3*k*_*B*_*T*, whereas the uncharged system has only a single, more extended basin. The first free energy minimum occurs for *G*_1_ *≈*0.6, *G*_2_ *≈*0.3, *R*_e2e_ *≈*0.5 nm and appears to correspond to a folded-over state that may evince intramolecular *β*-sheet characteristics at times, though based on the previous analysis, the formation of a true *β* sheet is comparatively rare. The second free energy minimum occurs for *G*_1_ 1. *≈*0, *G*_2_ *≈*0.3, *R*_e2e_ *≈*3.5 nm and appears to correspond to a primarily extended conformation, concomitant with the higher end to end distance and first gyration moment. When the system becomes uncharged, the free energy increases in the area of the first basin, but more collapsed states become more favorable overall, primarily those with slightly higher second moments. The barrier between states presumably occurs due to charge-charge repulsion between the lysines but evidently it is possible for a collapsed state to be stabilized, potentially due to the observed transient intramolecular hydrogen bonds (cf. Fig. 8a,d). Thus in the uncharged system we observe indiscriminate sampling of collapsed and extended states, whereas in the charged system we observe their separation by a modest free energy barrier.

Introducing multiple peptides to the system results in the resolution of more free energy minima separated by comparatively small barriers, primarily on the order of 2*k*_*b*_*T*, although the extent of the surface sampled is quite similar. We note also that the number of minima and the barrier heights tend to increase when the system becomes uncharged, rather than decrease as in the single-peptide case. We may rationalize this as the presence of larger aggregates in the uncharged system imposing heightened steric constraints on the peptides, which clearly has a greater effect in restricting the potential conformations than does the charge repulsion of the charged system. We note also the presence in both systems of a basin corresponding to extended states (*G*_1_ *>* 1.0, *R*_e2e_ *>* 3 nm) which may most easily participate in inter-molecular *β* sheets, although the extent of the basin is greater in the charged system and it appears to almost split into two in the uncharged system, potentially corresponding to differences in the arrangements of the two primary sites of *β* sheet (residues 3-6 versus 9-12, cf. snapshots in Fig. 9l, bottom right). One reason for the restriction of available conformational states may be that in the uncharged system the extended conformations are primarily stabilized by inter-molecular hydrogen bonding, whereas in the charged system, there is also an intra-molecular charge-charge repulsion component favoring more indiscriminate extended conformations. We note also 3-5 energy minima corresponding to collapsed states (*G*_1_ *<* 0.7, *G*_2_ *≈*0.35, *R*_e2e_ *<* 2.0 nm), including folded-over conformations like the single-peptide charged one in both charged and uncharged systems. These appear to be stabilized by both transient *α* helices and transient intra-molecular *β* sheets.

Overall, from these multi-peptide simulations we demonstrate the existence of an equilibrium of disparately-sized aggregates at neutral pH (charged system) that undergoes rapid aggregation when the pH is significantly increased (uncharged system) along with concomitant rapid formation of intermolecular *β* sheets in aggregates of size four and larger on similar timescales. It is possible that longer timescale rearrangements may lead to further enhancement in *β* sheet content, but it is unlikely to be by more than about 10%, according to experimental observations. Thus most *β* sheet formation occurs rapidly as part of aggregation. It appears to be seeded by transient aggregate formation in the charged system, which occurs primarily through C-terminal interactions, where the lower charge density is found, but charge neutralization leads to heightened N-terminal interactions, which appear to comprise the bulk of inter-molecular *β*sheet content in the high-pH system. Our observations are consistent with a comparatively simple picture in which the loss of charge-charge repulsion between monomers is the primary force driving both aggregation and conformational change in the system, as it allows for heightened peptide-peptide interaction that substantially restricts the conformational free energy landscape.

We note also the repeated motif of a folded-over collapsed hairpin-like structure, which in GL13K appears to be partially stabilized via intra-molecular hydrogen bonding and *β* sheet formation. In the A5 system, which experimentally demonstrates a slower aggregation rate, we observe a more consistent sampling of this folded-over state, potentially partially stabilized by a *β* bridge between ALA5 and LEU12 (cf. 4e). Since we observe the formation of *β*-sheet dimerization and transient larger aggregate formation even under charged conditions, we may explain A5’s slower aggregation by appealing to this in combination with the single-peptide simulations showing its heightened tendency to take on the collapsed hairpin-like state. In such a conformation, it would be unable to form dimers in the charged state; thus aggregation would become slower due to the lack of preformed dimers as nucleation sites. This provides further evidence for the picture of aggregation that we have presented and suggests that one avenue of future work might involve applying kinetic growth models to the non-equilibrium system, such as the Smoluchowski aggregation model, which has recently been generalized for small finite-size systems such as those observed in MD contexts ^[42]^.

Another point of interest is SER9. Replacement of SER9 with an ALA residue did not impact GL13K antibacterial activity but did impact its activity against the bacterial endotoxin LPS ^[17]^. In this context, we find it interesting to consider that SER9, in addition to being the only polar amino acid, is positioned centrally in the “hinge” of the collapsed conformation (cf. its participation in turn/bend secondary structure content, Figs 4, 7), as well as its comparatively elevated participation in both intraand intermolecular h-bonding. This suggests the possibility that in addition to the likelihood that SER9 controls interaction with LPS that the folded-over conformation may be relevant for LPS-GL13K interaction in specific. Since an increase in folded-over content can impact *β* sheet formation through removal of nucleation sites, it is of great interest to determine whether this conformation is indeed relevant to LPS binding and suggests that an important avenue of future work will involve simulations of these interactions.

## 3. Conclusion

We have performed the first *in silico* study of single-peptide and multi-peptide systems of the synthetic *β*-sheet forming peptide GL13K and its variants. We investigate systems with lysines charged, as they would be in neutral pH, such as in solution or in the proximity of a typical eukaryotic cell membrane, and with lysines uncharged (deprotonated), as they would be in high pH, such as in a high pH solution or possibly in the proximity of a typical negatively-charged prokaryotic cell membrane. We also simulate three alanine-scan variants where the fifth, the seventh, and the eleventh lysine are replaced with alanines, as well as two variants with the same amino acid composition but different sequences.

Our results match with experimental observations and allow us to go further by rationalizing the molecular-level underpinnings of those observations, localizing the stabilizing interactions along the peptide backbone, determining that the timescales for aggregation and primary *β*-sheet formation are commensurate, and providing a holistic illustration of the conformational free energy landscape and how it changes under different conditions that highlights the potential role of the equilibrium between collapsed and extended states.

An important area of future work includes investigation of GL13K in combination with different membrane variants. From this study, we have determined that at high-pH, it is sufficient to model aggregation according to deprotonation of the lysine residues via a simple neutral lysine model. It remains unclear, however, whether mitigation of charge-charge repulsion via electrostatic screening would have a similar effect. Thus, two possible pictures of aggregation at a bacterial membrane emerge, depending on the local pKa, which is notoriously difficult to model or measure. Either the local pKa is sufficiently modified to result in total deprotonation of lysine residues, or the electrostatic shielding of the positively charged lysines by the negatively charged bacterial membranes is sufficient to allow aggregation. Coarsegrained simulations under charged and uncharged conditions may be able to answer this question, as constant-pH simulations of fully atomistic membranes would be computationally prohibitive.

## 4. Experimental

We performed all-atom simulations of single and multi-peptide systems to investigate the effects of charge and sequence on conformation and aggregation of the peptide GL13K and related systems.

### 4.1 Single-peptide simulations

We conducted molecular dynamics simulations of GL13K, neutral GL13K, GL13KA5, GL13KA7, GL13KA11, GL13KR1, and neutral GL13KR1 using GROMACS version 2021.2 ^[43]^. Starting configurations for GL13K and GL13K-R1 were computed using Pepfold-3 ^[44]^ and employed as rough initial structures for all other peptides. GL13K, GL13K-neutral, GL13KA5, GL13K-A7, and GL13K-A11 were all started in the same configuration, and GL13K-R1 and GL13K-R1-neutral were both started in the same configuration.

Initial configurations were patched for C-terminal amidation and N-terminal protonated glycine using CHARMMGUI version 3.5 ^[45]^. We cap the termini rather than simulating the zwitterionic form to match previous experimental studies ^[21,25]^; it has been shown that differing terminal states can impact the conformations taken on by AMPs ^[46]^. We used the CHARMM36m force field version February 2021 in GROMACs ^[43]^, supplemented with an LSN residue (charge 0) parameterized by analogy with LYS (charge +1). The simulation environment is comprised of each AMP solvated using the TIP3P model for water in a cubic water box with sufficient counter-ions added to neutralize the system. The box dimensions extend 1.2 nm further than the dimensions of the AMP to prevent intermolecular interactions across periodic boundary conditions. Newton’s equations of motion were solved using a leap-frog algorithm ^[47]^. Van der Waals forces were truncated at 1.2 nm with a force switch smoothing function applied from 1.1-1.2 nm. Electrostatic interactions were calculated according to the smooth Particle Mesh Ewald method ^[48]^ with a real-space cutoff of 1.2 nm and a 0.12 nm Fourier grid spacing with a 4th order spline. The list of non-bonded (neighbour) particles was updated at least every ten steps and was generated with a Verlet scheme with a cutoff set to 1.2 nm. Hydrogen covalent bonds were constrained by LINCS to 4th order and a single correction iteration ^[49]^.

A steepest descent algorithm was used for energy minimization with a 0.01 step size and tolerance of *<* 1000.0 kJ/mol/nm to a maximum of 50 000 steps. A 100 ps NVT canonical equilibration was then conducted using a Nosé-Hoover thermostat ^[50,51]^, a bath with temperature 300 K, time constant 0.4 ps, and with protein and non-protein groups coupled separately. This was followed by a 100 ps NPT isothermal-isobaric equilibration using a Parrinello-Rahman barostat ^[52]^ with isotropic pressure coupling, a 1.0 bar reference pressure, a 2.0 ps time constant, and the compressibility of water set as 4.5 *×* 10^*−*5^ bar^*−*1^. The initial production run for each AMP was 30 ns with 2 fs time steps. The coordinates and energies were saved every 10 ps. We equilibrated each AMP for 200 ps followed by 500 ns of production.

### 4.2 Multi-peptide simulations

For both the GL13K and GL13K-neutral systems, we simulated eight peptides in a simulation box. Initial configurations for both charged and uncharged systems were taken from random configurations of the single-peptide simulation and solvated together. A summary of simulation time and setup can be found in tables S1 and S2. Simulations were run long enough to allow both cluster size and cluster *β*sheet content to reach steady-state values.

All MD simulations were performed with Lennard-Jones interaction cutoffs of 1.2 nm. Long-range electrostatic interactions were calculated using the particle-mesh Ewald method^[53]^ with conducting boundary conditions and a direct space cutoff of 1.2 nm. Simulations were performed in an isobaric-isothermal ensemble (NPT). The system pressure was maintained at 1 atmosphere with the Parrinello-Rahman barostat ^[54,55]^ using a 2.0 ps coupling time. The temperature was maintained at 300 K through a Nosé-Hoover thermostat ^[50,51]^ with 0.4 ps coupling frequency. All bonds between hydrogen and heavy atoms were constrained using the SHAKE algorithm ^[56]^ which permitted a 2.0 fs time step.

## Acknowledgements

This research was supported in part by Discovery Grant #RGPIN-2021-03470 from the National Sciences and Engineering Research Council of Canada. This research was enabled in part by support provided by Calcul Quebec (www.calculquebec.ca) and the Digital Research Alliance of Canada (https://alliancecan.ca). This research was undertaken, in part, thanks to funding from the Canada Research Chairs Program under grant number CRC-2020-00225. The authors thank Dr. Christine DeWolf and Dr. Deniz Meneksedag-Erol for useful discussions.

## Contributor Roles

Conceptualization: R.A.M., M.N.H.;

Methodology: R.A.M., L.N.W., M.N.H.;

Software: R.A.M., L.N.W., N.A.W., M.N.H.;

Validation: R.A.M., M.N.H.;

Formal analysis: N.A.W., M.N.H.;

Investigation: R.A.M., L.N.W., M.N.H.;

Resources: R.A.M.;

Data Curation: R.A.M., L.N.W., N.A.W., M.N.H.;

Writing - Original Draft: R.A.M., L.N.W., N.A.W., M.N.H.;

Writing - Review & Editing: R.A.M., N.A.W., M.N.H.;

Visualization: R.A.M., N.A.W., M.N.H.;

Supervision: R.A.M.;

Project administration: R.A.M. ;

Funding acquisition: R.A.M. ;

## Data availability

The authors confirm that the datasets containing molecular dynamics trajectories, structures, force fields and molecular dynamics input parameter files are publicly available at the following DOI: 10.5281*/zenodo*.10558960.

---

**Table S1.** Summary of simulation setup for single-peptide in solution.

|  | # replicas | sim. time (ns) | box type |
| --- | --- | --- | --- |
| R1-neutral | 5 | 500 | Cubic |
| R1 | 5 | 500 | Cubic |
| GL13K | 5 | 500 | Cubic |
| GL13K-neutral | 5 | 500 | Cubic |
| A5 | 5 | 500 | Cubic |
| A7 | 5 | 500 | Cubic |
| A11 | 5 | 500 | Cubic |

**Table S2.** Summary of simulation setup for multi-peptides in solution.

|  | # of peptides / replica | # of replicas | sim. time (ns) | box type |
| --- | --- | --- | --- | --- |
| Charged system | 8 | 5 | 1000 | Cubic |
| Uncharged system | 8 | 5 | 1500 | rhombic dodecahedron |

**Table S3.** Pairwise t-test of single peptide in solution on the radius of gyration data set. The upper triangle and lower triangle show p-values and T-values respectively.

|  | R1-neutral | R1 | GL13K | GL13K-neutral | A5 | A7 | A11 |
| --- | --- | --- | --- | --- | --- | --- | --- |
| R1-neutral | - | 0.18233 | 0.000007 | 0.0000014 | 0.00074 | 0.00003 | 0.00000006 |
| R1 | -1.46030 | - | 0.001373 | 0.008233 | 0.06858 | 0.00445 | 0.00183 |
| GL13K | -10.21573 | -4.78965 | - | 0.02379 | 0.01374 | 0.35347 | 0.16453 |
| GL13K-neutral | -12.67658 | -3.48707 | 2.78353 | - | 0.23103 | 0.18986 | 0.03862 |
| A5 | -5.28985 | -2.10335 | 3.14305 | 1.29625 | - | 0.0665 | 0.02913 |
| A7 | -8.41608 | -3.91546 | 0.98501 | -1.43261 | -2.12328 | - | 0.87404 |
| A11 | -18.96114 | -4.56676 | 1.53007 | -2.47147 | -2.65269 | 0.16368 | - |

**Figure S1.**
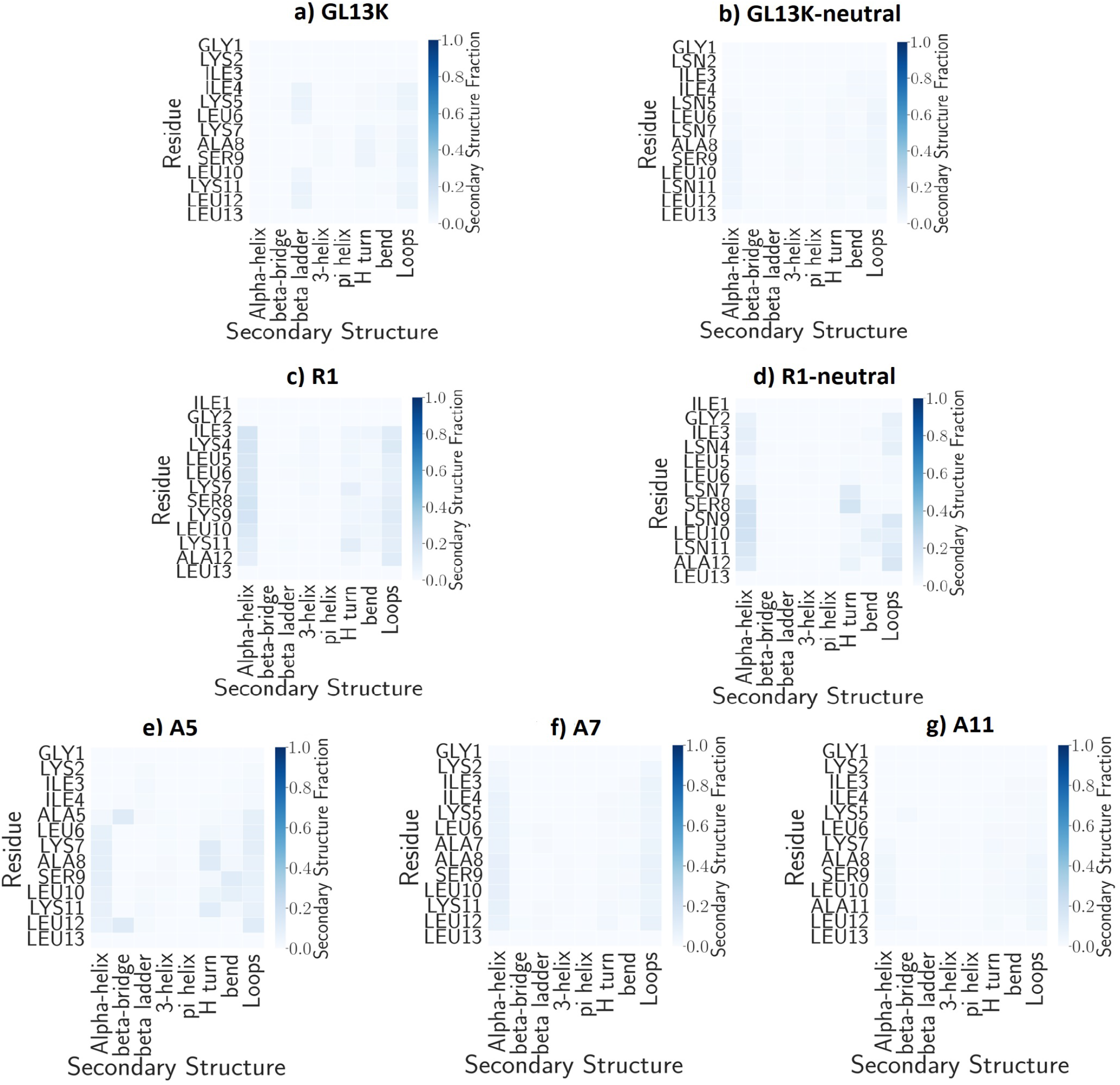
Average Secondary structure error for single-peptide systems in solution.

**Figure S2.**
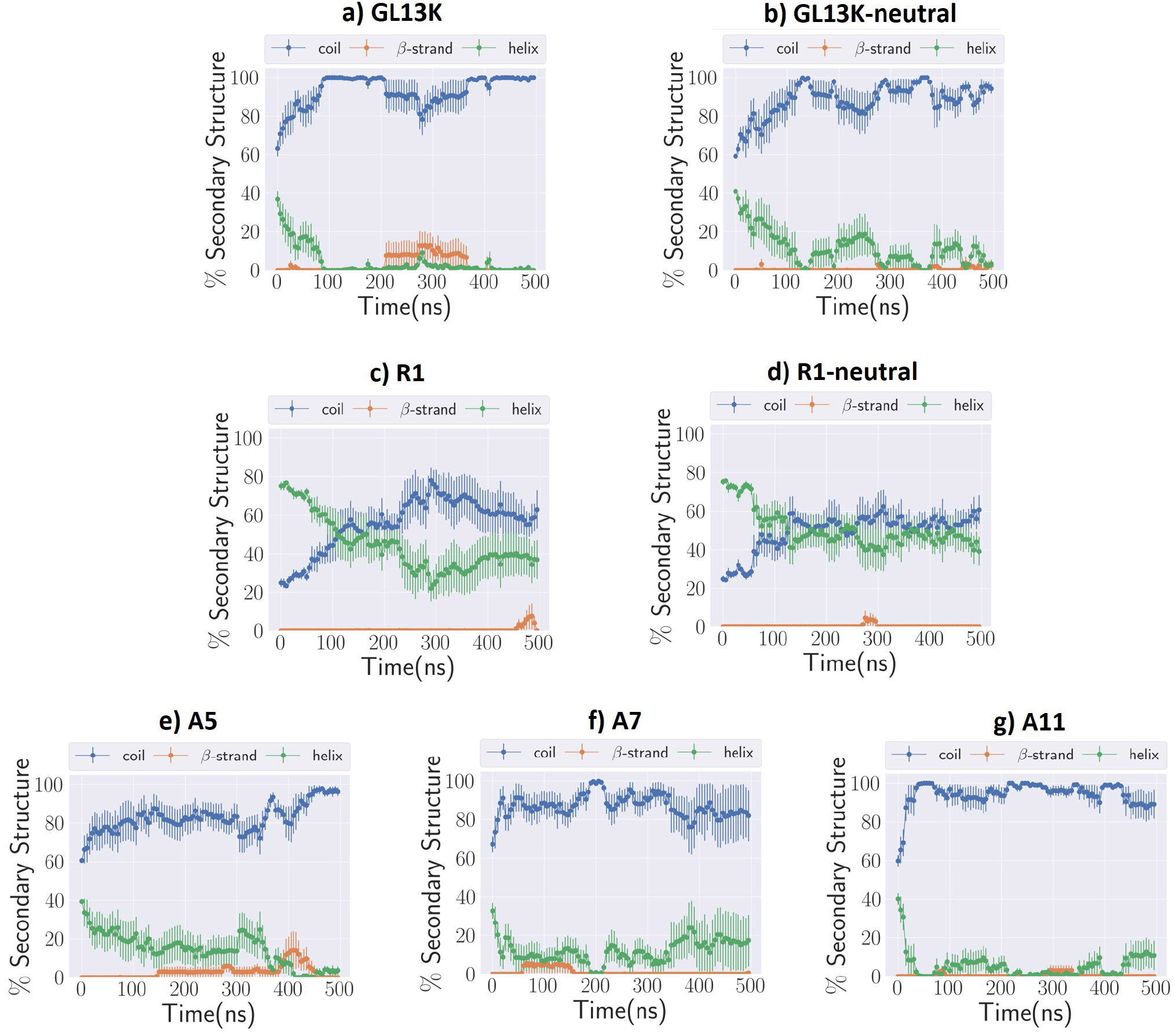
Secondary structure over time for single-peptide simulations of (a) GL13K, (b) GL13K-neutral, (c) R1, (d), R1-neutral, (e) A5, (f) A7, (g) A11. We show the average and standard error of five independent replicas, and we focus on helix, random coil, and *β*-strand content.

**Figure S3.**
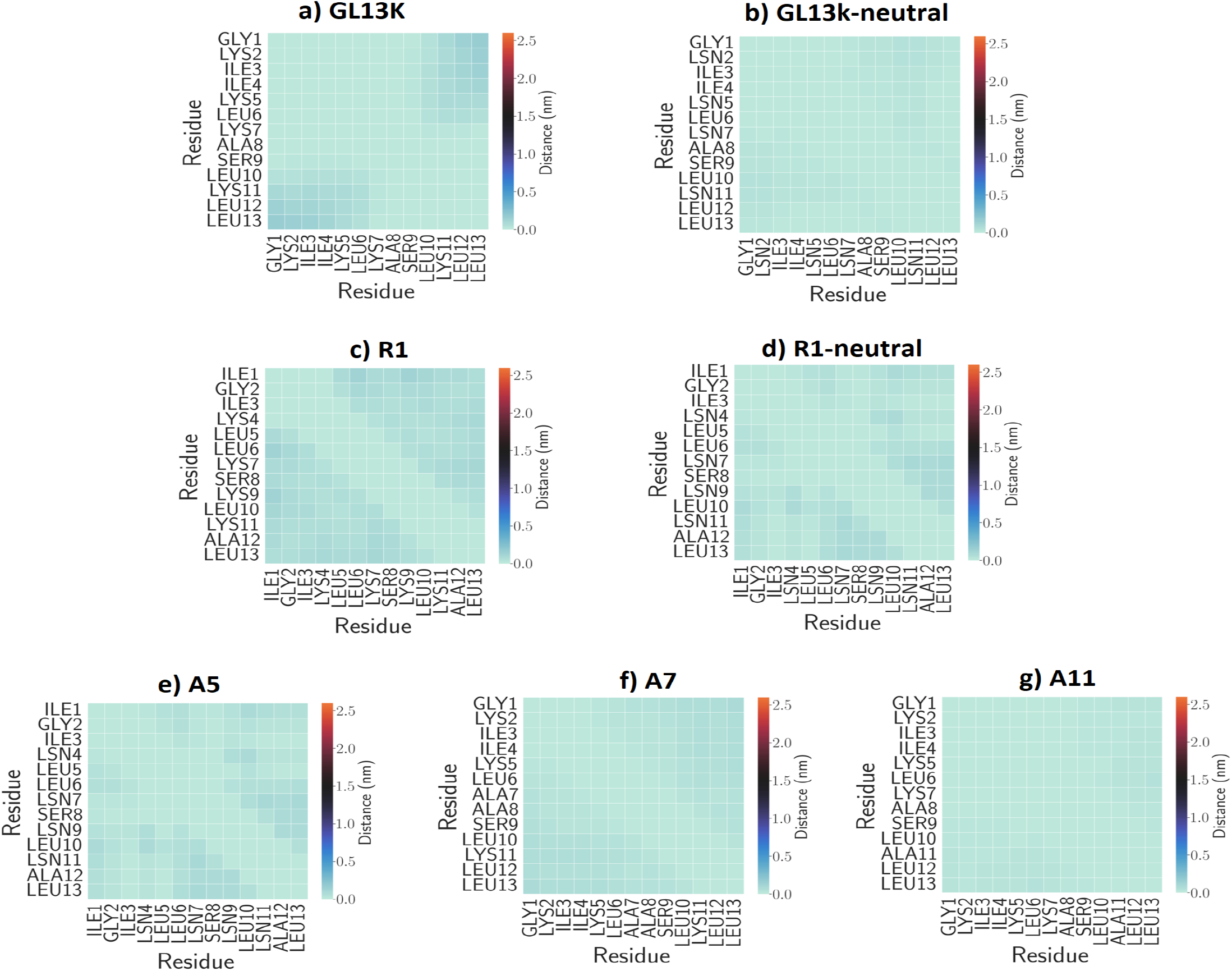
Average contact map error of single-peptide systems in solution

**Figure S4.**
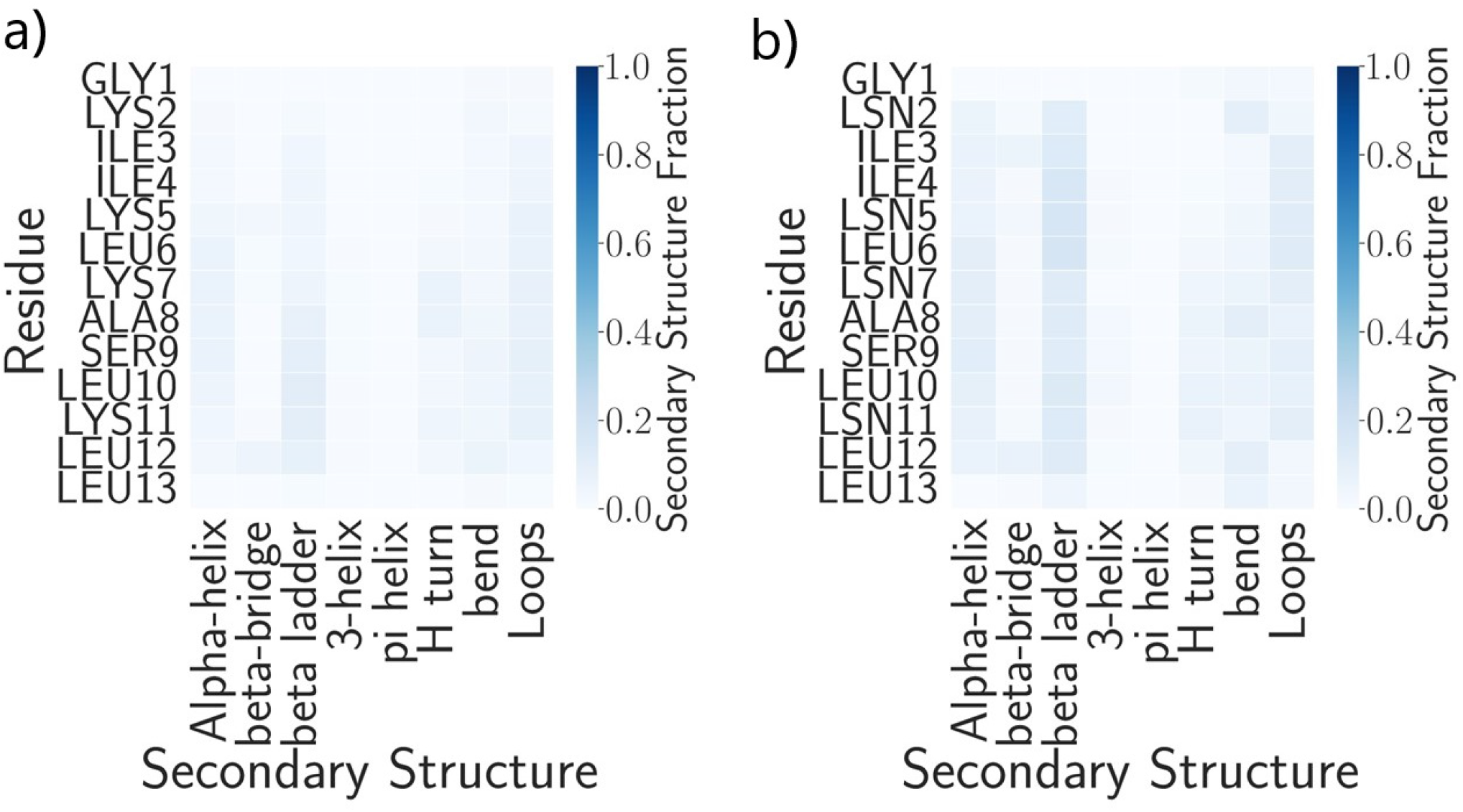
The error of average secondary structure per residue of multi-peptide systems: (a) charged system; (b) uncharged system.

